# Objective Extraction of Evoked Event-related Oscillations from Time-frequency Representation of Event-related Potentials

**DOI:** 10.1101/2020.05.17.100511

**Authors:** Guanghui Zhang, Tapani Ristaniemi, Fengyu Cong

## Abstract

Evoked event-related oscillations (EROs) have been widely used to explore the mechanisms of brain activities for both normal people and neuropsychiatric disease patients. The selection of regions of evoked EROs tends to be subjectively based on the previous studies and the visual inspection of grand averaged time-frequency representations (TFRs) which causes some missing or redundant information. Meanwhile, the evoked EROs cannot be fully extracted via the conventional time-frequency analysis (TFA) method because they are sometimes overlapped with each other or with artifacts in time, frequency, and space domains to some extent. Hence, these shortcomings may pose some challenges to investigate the related neuronal processes. A data-driven approach was introduced to fill the gaps as below: extracting the temporal and spatial components of interest simultaneously by principal component analysis and Promax rotation and projecting them to the electrode field to correct their variance and polarity indeterminacy, calculating the TFRs of the back-projected components, and determining the regions of interest objectively using the edge detection algorithm. We performed this novel approach and the conventional TFA method in analyzing both a synthetic dataset and an actual ERP dataset in a two-factor simple gambling paradigm of waiting time (short/long) and feedback (loss/gain) separately. Synthetic dataset results indicated that N2-theta and P3-delta oscillations were detected using the proposed approach, but, by comparison, only one oscillation was obtained via the conventional TFA method. Furthermore, the actual ERP dataset results of P3-delta for our approach revealed that it was sensitive to the waiting time (which also was found in the previous reports) but not for that of the conventional TFA method. This study manifested that the proposed approach can objectively extract evoked EROs, which allows a better understanding of the modulations of the oscillatory responses.

## 1. Introduction

EEG is widely used in neuroscience to evaluate the temporal, spectral, and spatial dynamics of cognitive processes. One typical technique is the event-related potential (ERP), which is computed by averaging multi-trial EEG data, and the other one is the evoked event-related oscillations (EROs) in the time-frequency domain or the spectral domain based on the ERPs (Başar et al., 2001). The evoked EROs have been widely used for investigating the distinctions between the related cognitive functions in normal and neuropsychiatric disordered subjects (Başar et al., 2016, 2013). Several approaches have been developed for the evoked EROs, such as digital filtering (like 4-8Hz for theta band), power spectral density-based spectral analysis, and time-frequency analysis (TFA) (Güntekin & Başar, 2016). The underlying ideas to calculate the evoked EROs by digital filtering technique and spectral analysis are similar. For the digital filtering method, the EROs are obtained by using a bandpass filter to filter the signals with the demanded frequency bands. The main drawback of such an approach is that it is difficult to see how the EROs change with frequencies in each time point for the filtered signals. TFA can overcome this obstacle, allowing examination of EROs in temporal and spectral domains simultaneously. The regions of evoked EROs of interest were usually determined by a certain time window and a frequency range according to the previous studies and the visual inspection of grand averaged time-frequency representation (TFR) distributions - in other words, they were subjectively chosen by experimenters (Ergen et al., 2014; Lally et al., 2014; Jones et al., 2006; Sandre & Weinberg, 2019; Zhang et al., 2019; Her et al., 2019). This technique is called as conventional rectangle method in this study due to the rectangle-like shape of this certain region. Most noticeably, the shapes of evoked EROs are more like a waterdrop than a rectangle and they are not the same as each other for TFR distributions of all conditions of ERP signals. Generally, the same certain region is used for all conditions when using this approach to determine the regions of interest. As a result, some useful information is neglected if the marked region is smaller than the shape of evoked ERO or unrelated information is involved when the marked region is larger than the real boundary of evoked ERO see the regions were marked by the conventional rectangle method in Figs. 6 - 9 **(B)**. The other thing should be considered, as shown in Figs. 3 and 4 corresponding to the waveform of a mixed synthetic signal, two ERPs (i.e., N2 and P3, they are called as ‘ERP’, and during the temporal principal component analysis (t-PCA) and rotation steps, we use the notation of ‘component(s)’ to represent the rotated principal components and the projected ones in this study) are used for further analysis and they are overlapped in time, space and frequency domains to some degree. While only one evoked ERO is recognized in the TFR distribution of the mixed-signal which is obtained by the conventional TFA method (as shown in Figs. 3 and 4 **(B)** the associated TFR distribution of the mixed-signal). This can be also confirmed in the actual ERP signals (Barry, 2009; Ergen et al., 2014; Jones et al., 2006). In their studies, the number of visible evoked EROs identified from the grand averaged TFR was smaller than those of the analyzed ERPs. Thus, it remained challenges that some stages of cognitive processes cannot be completely interpreted.

To address these gaps, we proposed an approach to objectively extract evoked EROs (as shown in Fig. 1). More specifically, t-PCA and Promax rotation were first conducted to extract the temporal and spatial components, and the selected components of interest were projected to the electrode field to correct their variances and polarities indeterminacy. Then, we applied a complex morlet continuous wavelet transformation to calculate the TFRs of the back-projection. Finally, the edge detection algorithm based on Canny detector was introduced to determine the specific time and frequency positions of evoked ERO of each condition from the associated TFR distribution for statistical analysis. Additionally, the homogeneity of topography across all participants was applied to evaluate the stability of ERPs/components/evoked EROs.

**Fig. 1.**
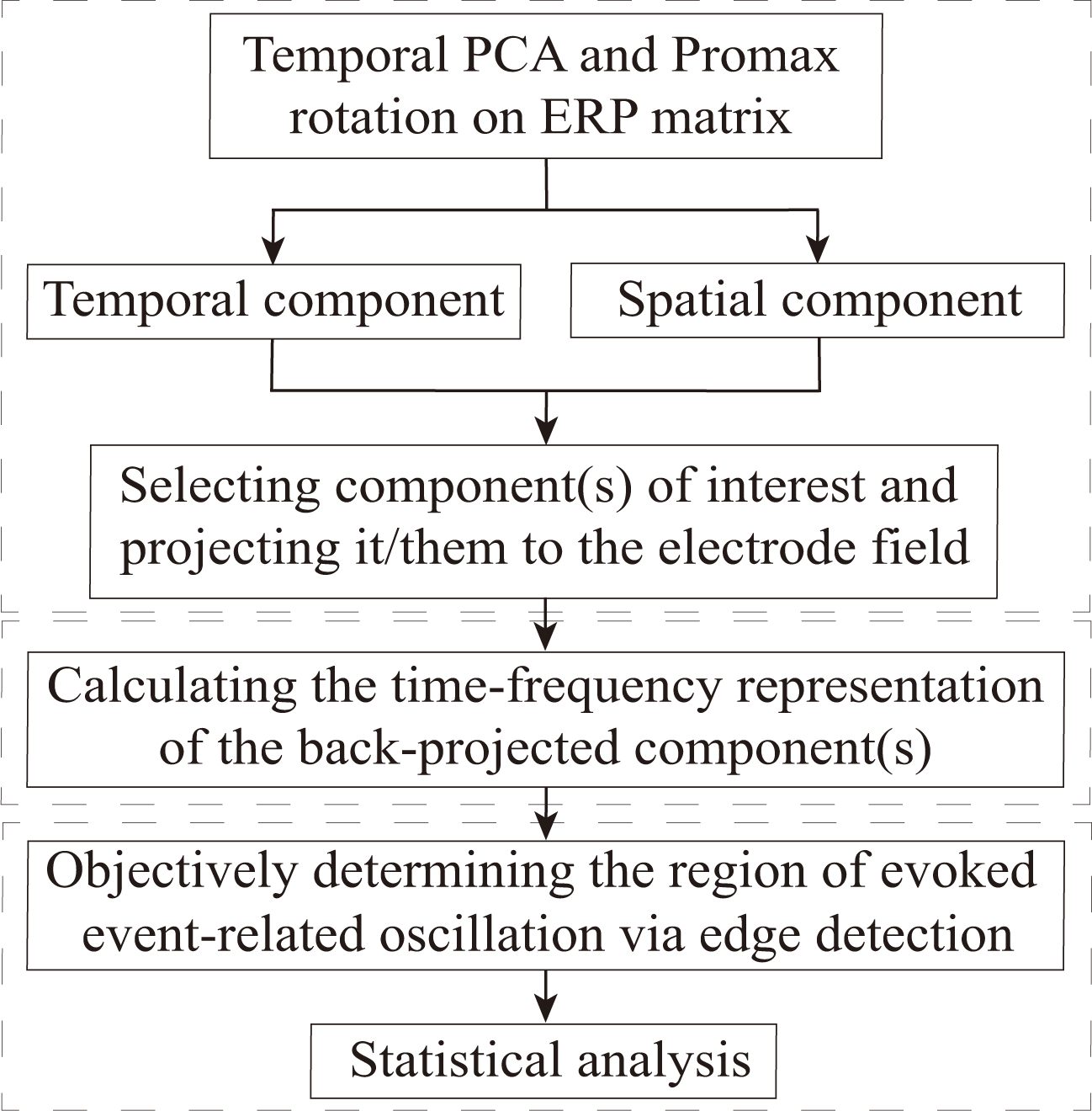
The diagram for extracting evoked event-related oscillations (EROs) from ERP datasets using the proposed method. First, exploring the temporal-spatial components of interest using temporal principal component analysis (t-PCA) and Promax rotation and projecting them to the electrode field. Second, transforming the projection of the components of interest into time-frequency representations (TFRs) via complex morlet continuous wavelet transformation. Third, determining the time and frequency positions of the evoked eventrelated oscillation of interest objectively using edge detection algorithm for statistical analysis.

There were two assumptions in this study. On the one hand, we presumed that the proposed approach can objectively and efficiently extract the temporal and spatial components in the time and space domains and temporal-spectral feature of evoked EROs in the time-frequency domain from the mixed-signal but do not change the associated temporal, spatial and TFRs characteristics. On the other hand, we also supposed that the statistical results of the proposed approach were not subject to the experimenters. To demonstrate these two aspects, the waveforms/topographies/TFRs of the source, mixed, and extracted simulation signals were displayed and the correlation coefficients (CCs) between any two of these of the three signals were calculated in the first step. Furthermore, the conventional TFA method and the proposed approach were used to obtain the evoked P3-delta and N2-theta oscillations from the actual dataset separately. For the determination of the region for the evoked ERO of interest, it was achieved by the conventional rectangle method and edge detection algorithm separately, and a comparison was made between the statistical results of these two methods. The codes for the proposed method of this investigation can be found from the link: http://www.escience.cn/people/cong/Toolbox.html.

## 2. Methods

### 2.1. EEG and synthetic datasets collection and analysis

#### 2.1.1. Synthetic dataset

The synthetic signal was generated with Dipole-Simulator (BESA Tool version 3.3.0.2). There were four simulated ERPs (N1, P2, N2, and P3) whose maximum amplitudes were measured at Fz, CPz, FCz, and Cz respectively. The details of their associated waveforms, topographic maps in the time domain, and TFR distributions can be found in Fig. 2. Meanwhile, we also displayed CCs between any two of the time-course/topographic maps/TFRs of the four original sources and their mixture to show the degree of overlap among them and how much the four sources contribute to the original mixed-signal (see the last row in Fig. 2). The duration of the signal was 1000ms (from −200ms to 800ms). In addition, the sampling rate was 150Hz and the background RMS noise level was set to 0 *µV*. In this study, N2 and P3 were considered as the interested ERPs and the others were concomitant ones. The maximum peak for N2 and P3 located in 260-390ms and 370-580ms separately. In order to simulate the signals as close to the actual ERP signals as possible, the variations were set in latency and amplitude of P3 and N2 of the original mixed-signal respectively, which was applied to simulate the single-trial dataset (David et al., 2006). Following this idea, the 11-set data were then simulated. The white Gaussian noise was added to the mixed 11-set signals (as shown in Figs.3 and 4, the mixed-signal plays the role of the preprocessing ERPs with the consideration of a real ERP dataset) using Matlab function-*awgn*, in which the signal-noise-ratio (SNR) was set to 10dB.

**Fig. 2.**
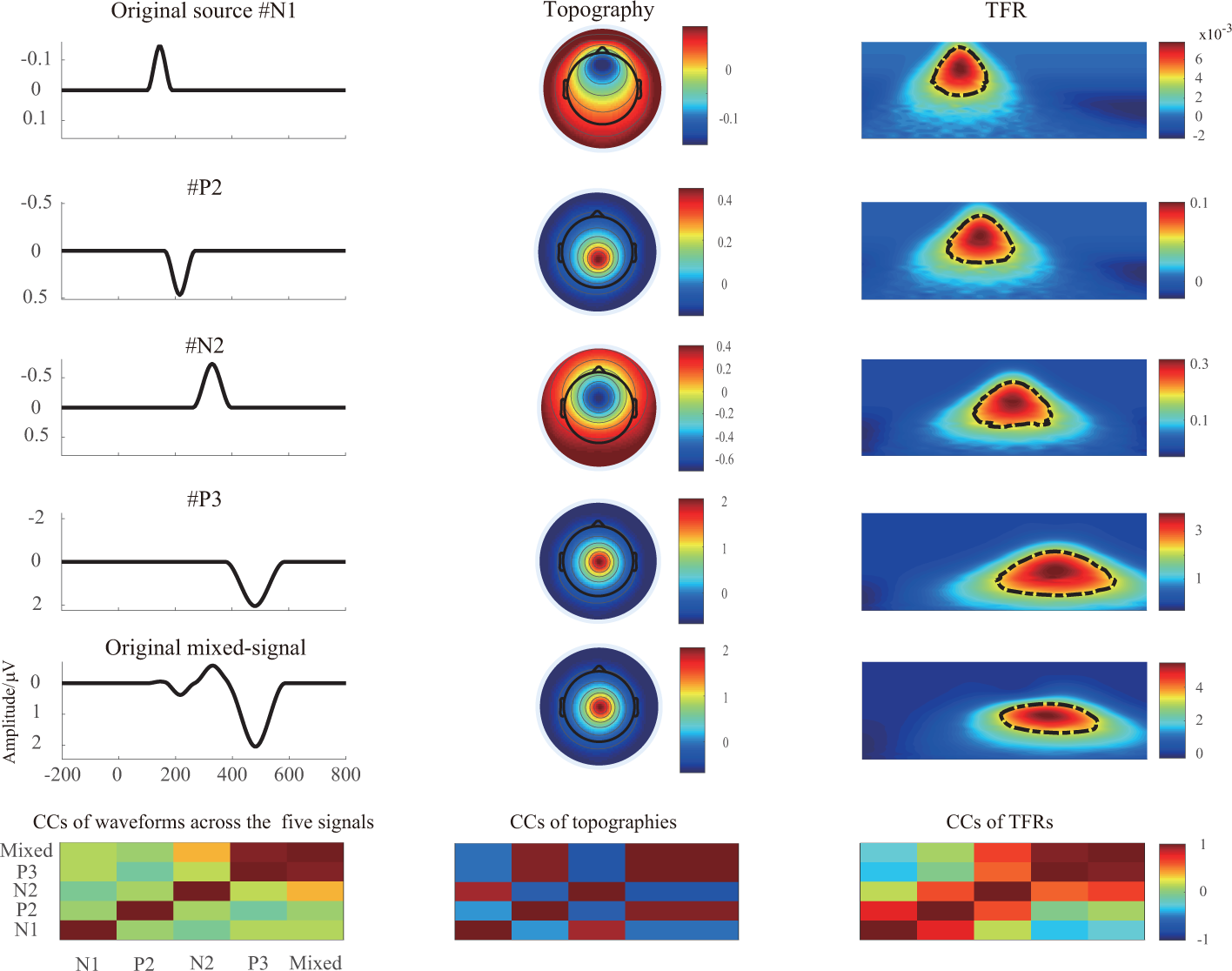
The waveforms, associated topographies, and time-frequency representations (TFRs) for the original source N1 (Fz), P2 (CPz), N2 (FCz), P3 (Cz), and mixed-signal (Cz) respectively (the first five rows). The last row represents the correlation coefficients (CCs) of waveforms/topographies/TFRs among all five signals. The 11 sets of simulation signals were generated from these sources based on setting the variations of amplitude and latency for N2/P3.

#### 2.1.2. ERP dataset

Twenty-one undergraduate and graduate students were recruited to participate as paid volunteers. Nine were females and twelve were males (mean age: 20.95 years old). All the participants were right-handed, with normal or corrected to normal visual acuity and they did not know or see the experimental paradigm before the experiment. The details of the experiment materials and the paradigm can be found in this research (Wang et al., 2014). EEG recordings at 64 locations were collected according to the standard 10-20 system (Brain Products GmbH, Gilching, Germany). The EEG data were referenced online against the left and right mastoids. Meanwhile, a vertical electrooculogram (EOG) was obtained below the left eye, and the horizontal electrooculogram was obtained at the outer canthus of the right eye. All impedances were less than 10kΩ for each sensor of every participant. The EEG and EOG for each participant were recorded with a 500Hz sampling rate and the data were filtered between 0.01 and 100Hz using a band-pass filter.

#### 2.1.3. Data preprocessing and analysis

##### Synthetic dataset

According to the recommendations in our previous study (Cong et al., 2015) and the frequency band of components of interest, the synthetic data were first filtered using wavelet filter (Cong et al., 2012, 2015) with these parameters - the number of levels for decomposition was 8; the selected mother wavelet was ‘*rbio*6.8’; the detail coefficients of the number of levels at 4, 5, 6, 7, and 8 were chosen for signal reconstruction. The TFRs were calculated via wavelet transformation for the source, mixed, and the extracted signals separately. Meanwhile, aiming at obtaining better time-resolution and frequency-resolution of TFRs, the center frequency and bandwidth were set as 1 respectively to define the mother wavelet as applied in our previous study (Zhang et al., 2020), and the frequency range of interest was from 1 to 15Hz with 40 frequency bins in nonlinear distribution. For each frequency layer, the power values were baseline corrected by subtracting the mean power of the baseline (200 ms before the stimulus onset) for each point using the subtraction approach (Benvenuti et al., 2017; Hu et al., 2014; Peng et al., 2019).

To reveal that the proposed approach can efficiently extract the evoked ERO from the ERP waveforms in the time domain without changing their temporal/spatial/TFR properties, the related waveforms, the associated topographies, and TFRs are plotted for the source, extracted, and mixed signals (see Figs. 3 and 4). Furthermore, CCs between any two of the waveforms/topographies/TFRs of the source, mixed, and extracted signals are computed to further prove this assumption as illustrated in Fig. 5.

**Fig. 3.**
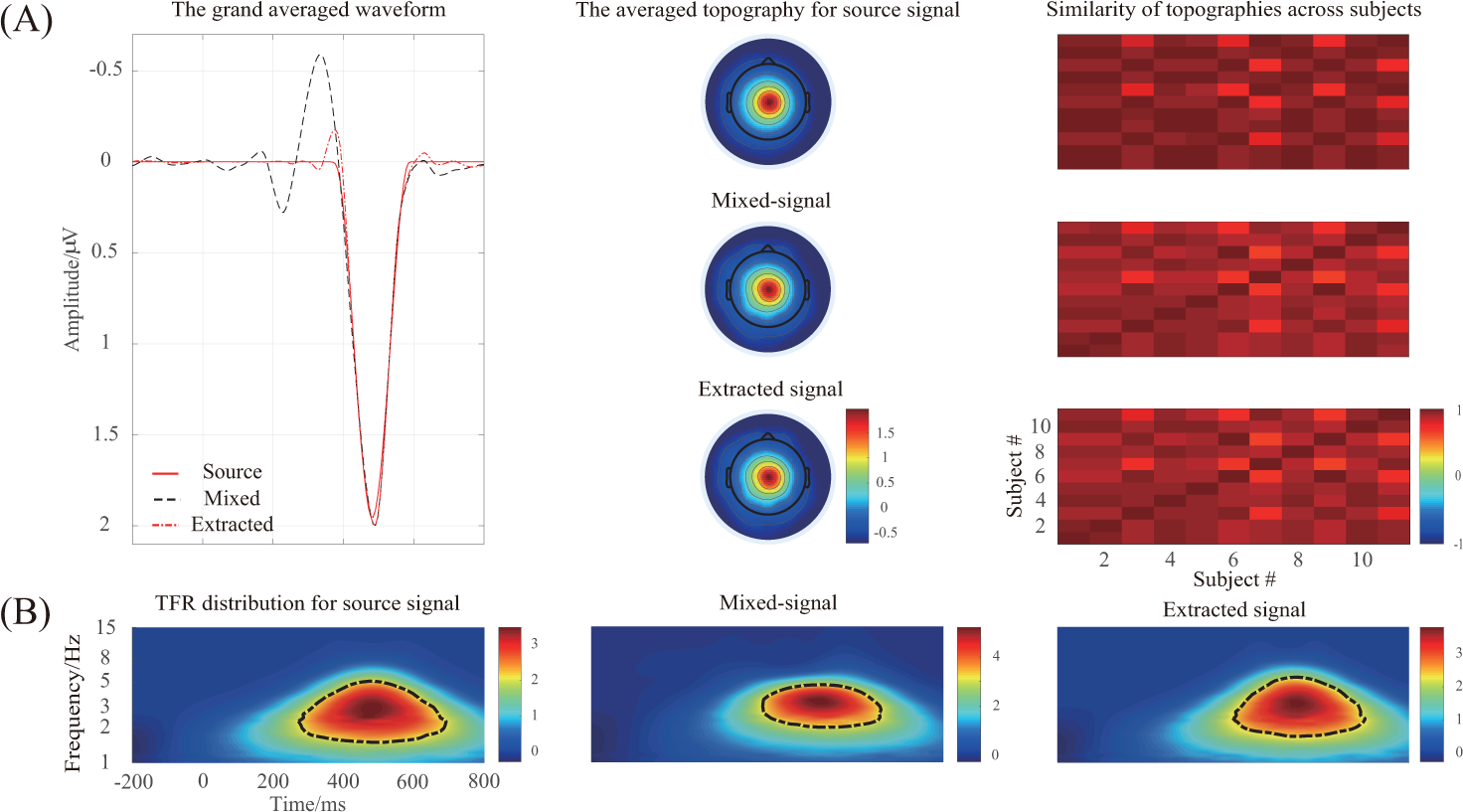
(A) The grand averaged waveform at Cz electrode, topography(time window was 450-550ms), and similarity of topographies among all subjects of the source/mixed/extracted P3 for the synthetic dataset. (B) The grand averaged time-frequency representation (TFR) at Cz of the source, mixed and extracted P3-delta separately. The mixed-signal plays the role of the preprocessing ERPs with the consideration of a real ERP dataset.

**Fig. 4.**
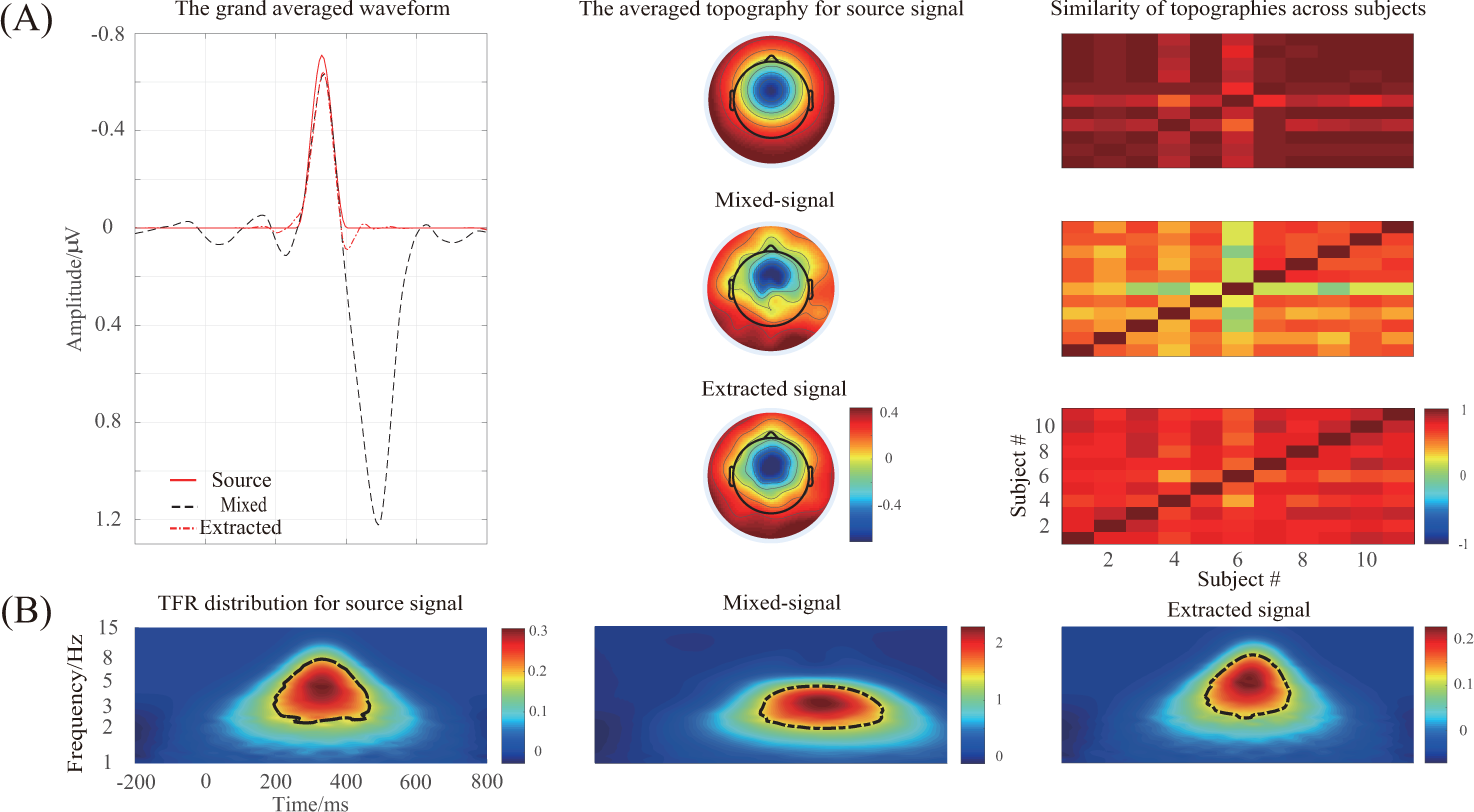
(A) The grand averaged waveform at FCz electrode, topography (it was measured within the time window 300-400ms), and similarity of topographies among subjects of the source/mixed/extracted N2 for the synthetic dataset. (B) The grand averaged time-frequency representation (TFR) of N2-theta for the three signals. The mixed-signal plays the role of the preprocessing ERPs with the consideration of a real ERP dataset.

**Fig. 5.**
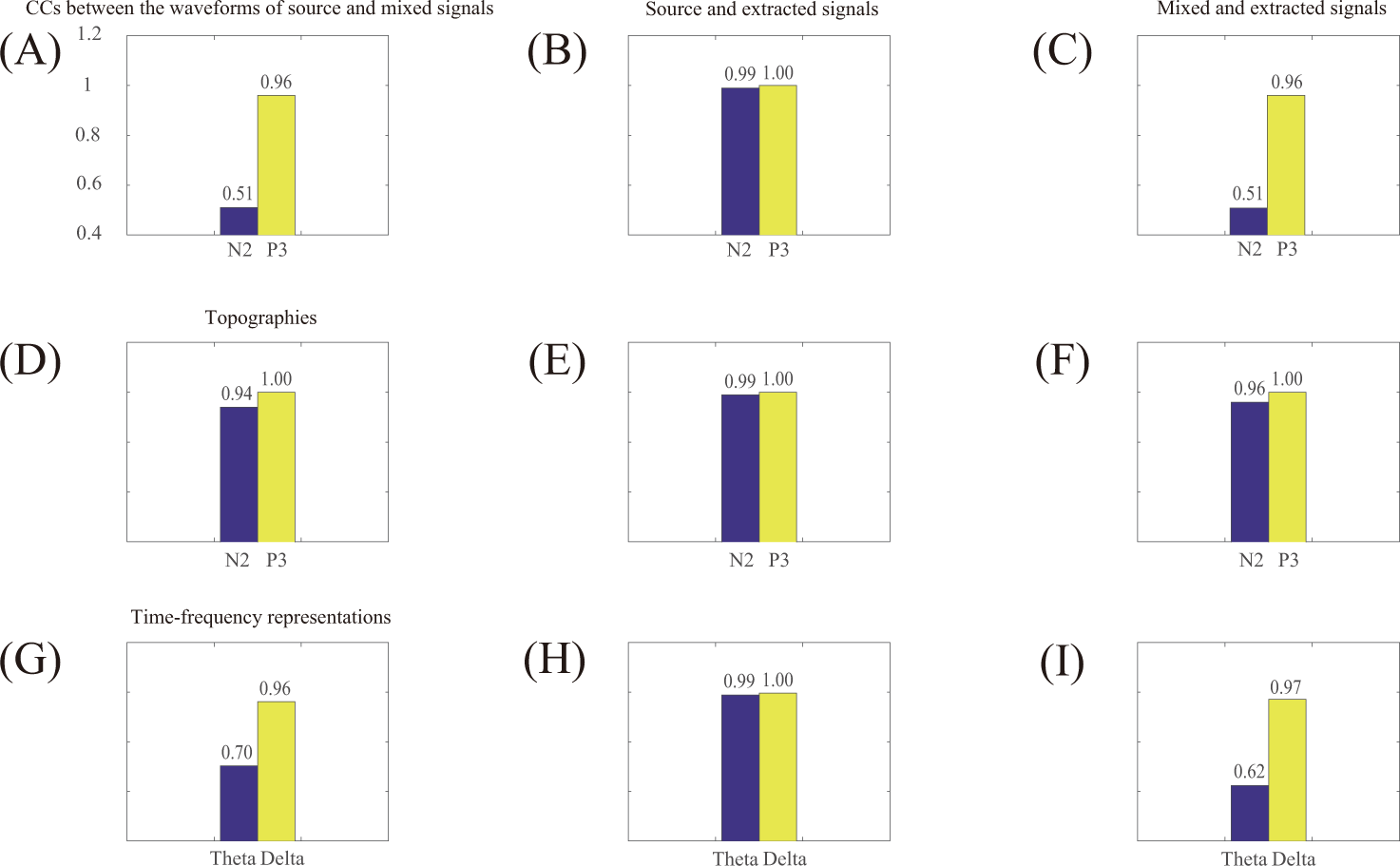
The correlation coefficients (CCs) between any two waveforms/topographies/timefrequency representations (TFRs) of the source, mixed, and extracted N2 (theta)/P3 (delta) for the synthetic dataset.

##### Actual dataset

The EEG signals were filtered offline using a notch FIR filter with 45-55Hz, a high-pass FIR filter of 0.1Hz, and a low pass FIR filter with 30Hz. Then, the filtered continuous recordings were segmented from 200ms before the stimulus onset to 1000ms after the stimulus onset. Epochs/trials whose maximum magnitude exceeded 100*µV* were excluded and the remaining epochs were baseline corrected. Next, the multi-trial dataset was averaged across every condition of each participant and the averaged datasets were then filtered by the wavelet filter to improve the SNR. Combining the recommendations mentioned in our previous study (Cong et al., 2015) and the frequency bands of components, the parameters for wavelet filter were finally set as following: the number of levels for decomposition was 9; the selected mother wavelet was ‘*rbio*6.8’; the number of levels for reconstruction was 5, 6, 7, 8, and 9.

The parameters for the conventional TFA method were the same as the applications in the synthetic dataset analysis. Two evoked EROs (P3-delta and N2-theta) were analyzed via the conventional TFA method and the proposed approach separately. Two-way repeated-measurement-ANOVA (rm-ANOVA) with waiting time (short/long) and feedback valence (loss/gain) as within-subject factors was used for statistical analysis which was performed in the SPSS 21.0 environment. The correction of the number of degrees of freedom would be carried out by the Greenhouse-Geisser method if necessary.

1. Conventional time-frequency analysis method. The conventional rectangle method and edge detection algorithm were used to obtain the regions of delta and theta oscillations respectively. When the conventional rectangle method was applied to determine the region, two regions were used for the delta and theta oscillations separately. For the N2-theta mean power was obtained by calculating the mean value of every condition across subjects with the regions (100-300ms, 3-7Hz; 50-350ms, 3-7Hz as shown in Fig. 6 **(B)**, they correspond to the black dotted and red solid rectangles respectively) at Fz, FCz, and Cz electrodes. And the mean values of two different regions (200-600ms,1-3Hz; 300-600ms, 1-4Hz as shown in Fig. 7 **(B)**, they were marked using the black dotted and red solid rectangles respectively) of every subject were measured for P3-delta power at Fz, FCz, Cz, CPz, and Pz electrodes respectively.
2. Proposed approach. The region of ERO under each condition was marked using the two methods as mentioned above (for the conventional rectangle method, N2-theta: 4-8Hz, 150-300ms see the red solid rectangle in Fig. 8 **(B)**; P3-delta: 1-3Hz, 200-600ms see the red solid rectangle in Fig. 9 **(B)**) respectively for further statistical analysis. As described in the conventional TFA method, the power of P3-delta and N2-theta oscillations were also measured at the same electrodes respectively.

**Fig. 6.**
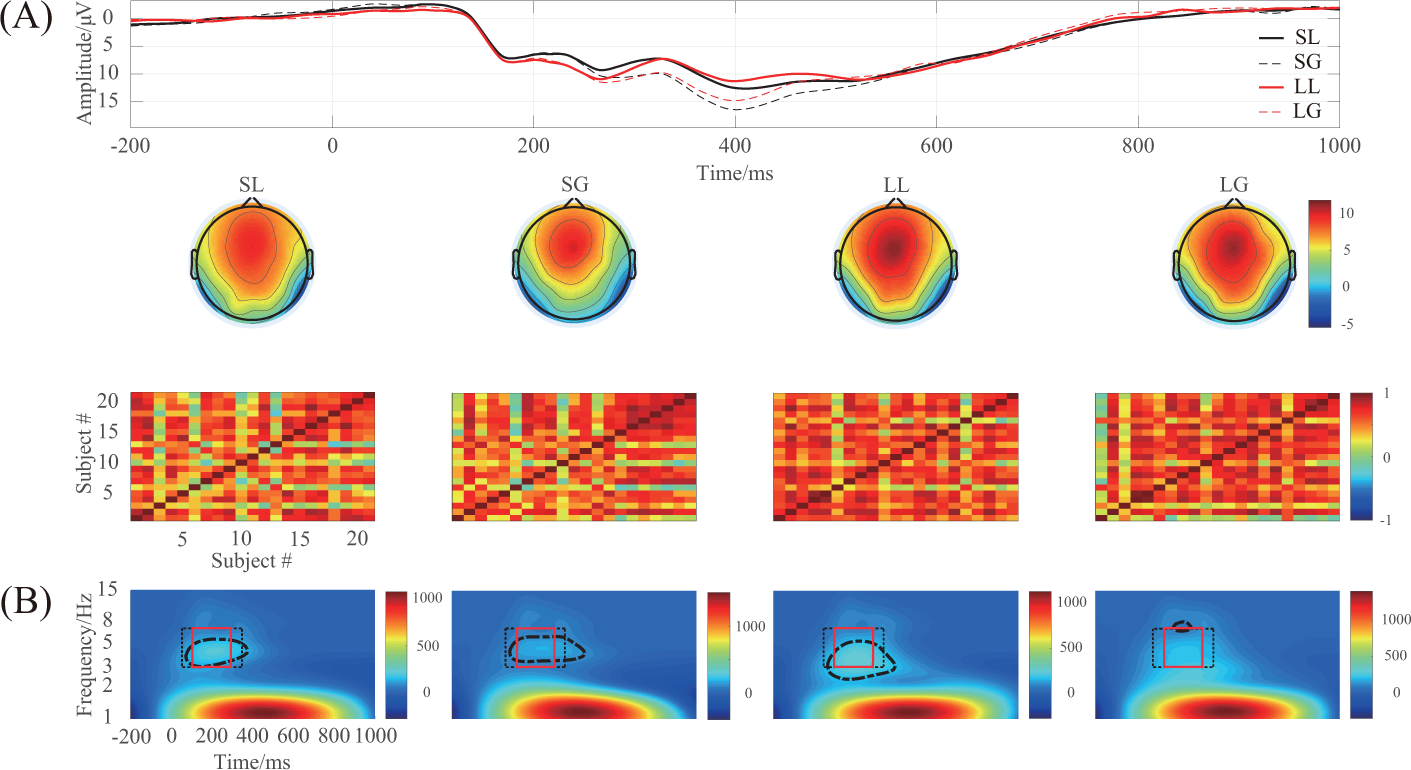
(A) The grand averaged waveform (at Fz, FCz, and Cz electrodes), topography (time window: 180-240ms), and similarity of topographies across participants of each condition for the filtered real signal. (B) The associated grand averaged time-frequency representation (TFR) of every condition. The region of evoked ERO of each condition is determined via the edge detection algorithm and the conventional rectangle method (for the black dotted rectangle, the time window is 50-350ms and frequency range is 3-7Hz; the red solid rectangle: 100-300ms and 3-7Hz) separately. SL: loss condition under short waiting time; SG: gain condition under short waiting time; LL: loss of long waiting time; LG: gain of long waiting time.

**Fig. 7.**
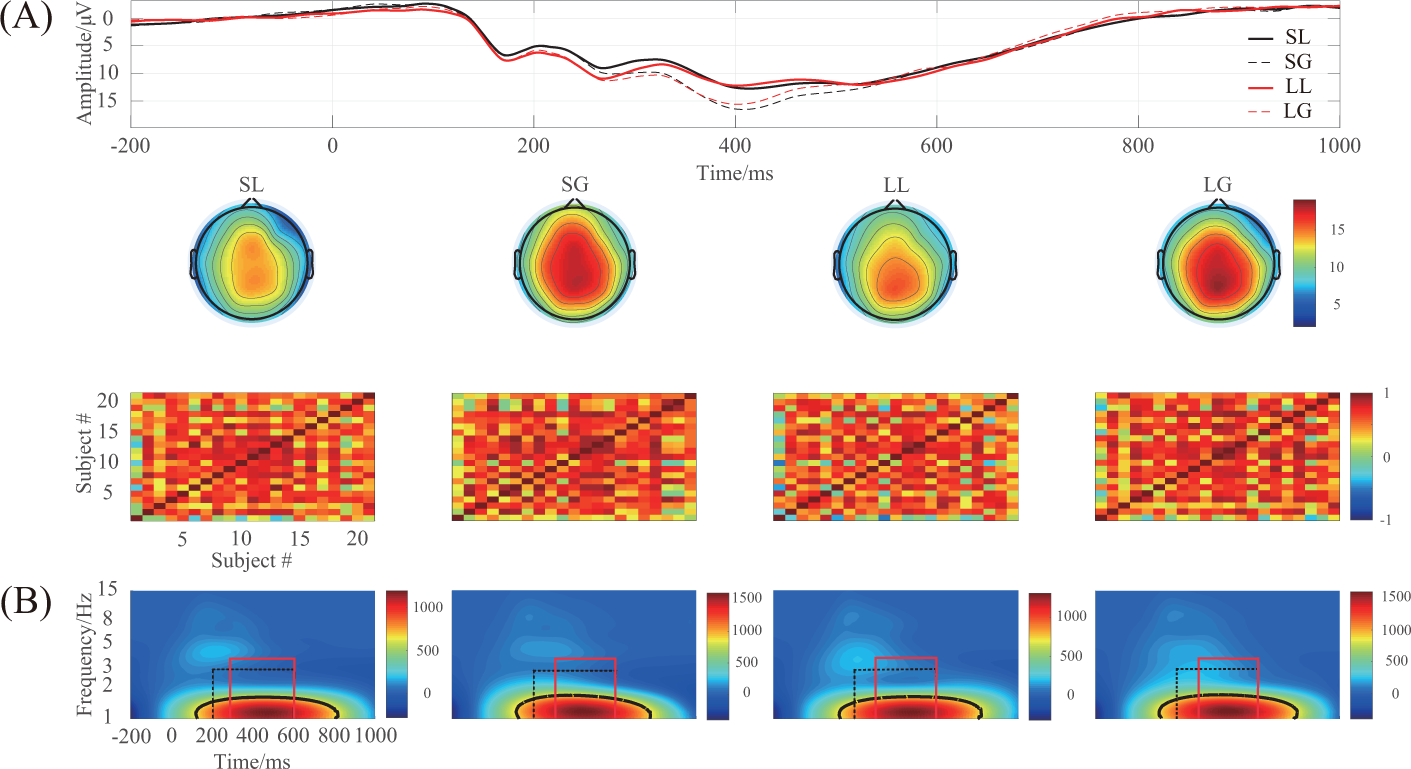
(A) The grand averaged waveform (at Fz, FCz, Cz, CPz, and Pz electrodes), topography (time window: 300-600ms), and similarity of topographies across participants of each condition for the filtered real signal. (B) The associated grand averaged time-frequency representation (TFR) of every condition. The region of evoked ERO of each condition is determined via the edge detection algorithm and the conventional rectangle method (for the black dotted rectangle, the time window is 200-600ms, and frequency range is 1-3Hz; the red solid rectangle: 300-600ms and 1-4Hz) separately.

**Fig. 8.**
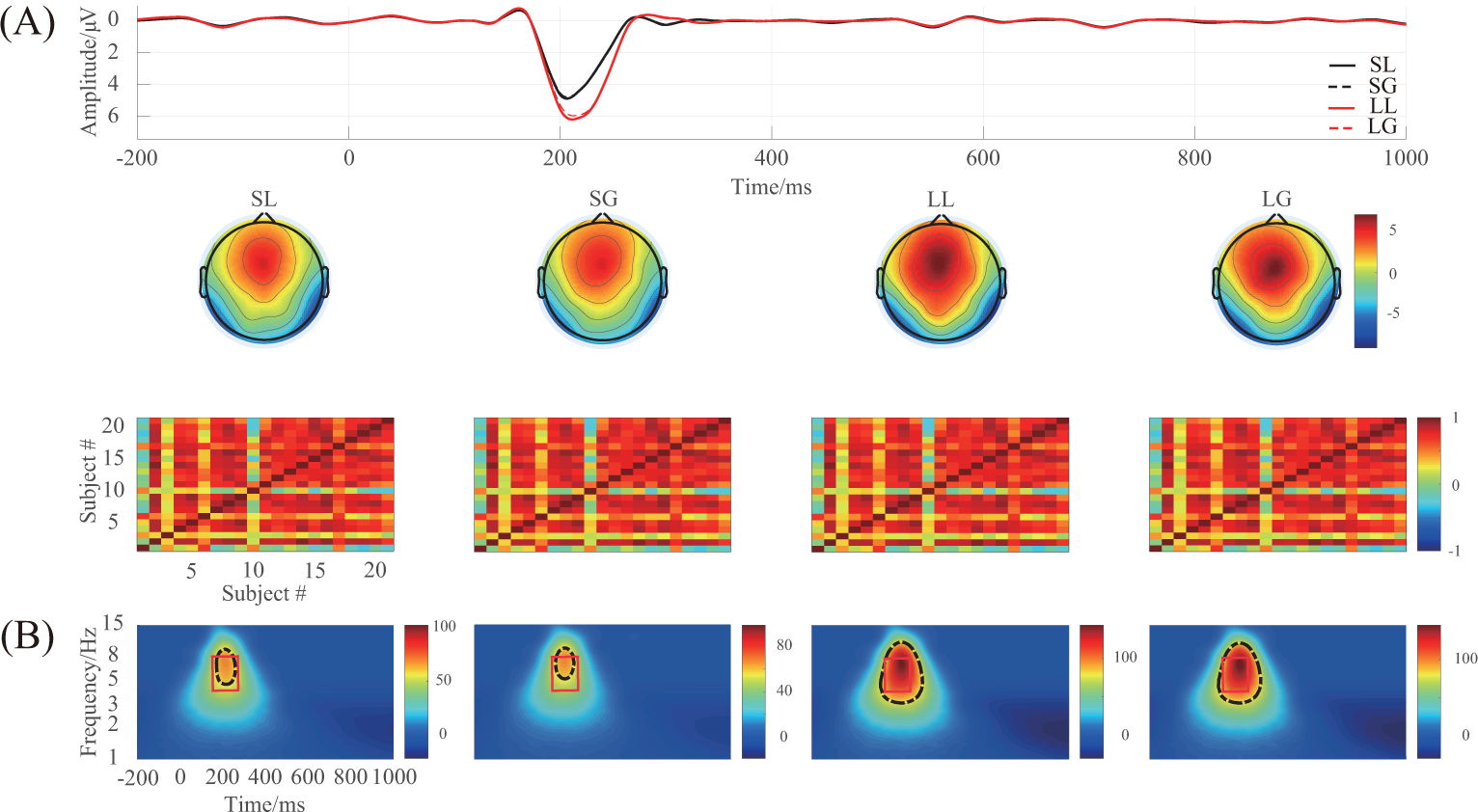
(A) The projected waveform (at Fz, FCz, and Cz electrodes), topography (180-240ms), and similarity of topographies across participants of every condition for N2 which were extracted from the real mixed-signal using t-PCA and Promax rotation. (B) The associated grand averaged time-frequency representation (TFR) of every condition for N2-theta oscillation. The red solid rectangle (the time window is 150-300ms and the frequency range is 4-8Hz) for every condition was marked using the conventional rectangle method, the other was gained by the edge detection algorithm.

**Fig. 9.**
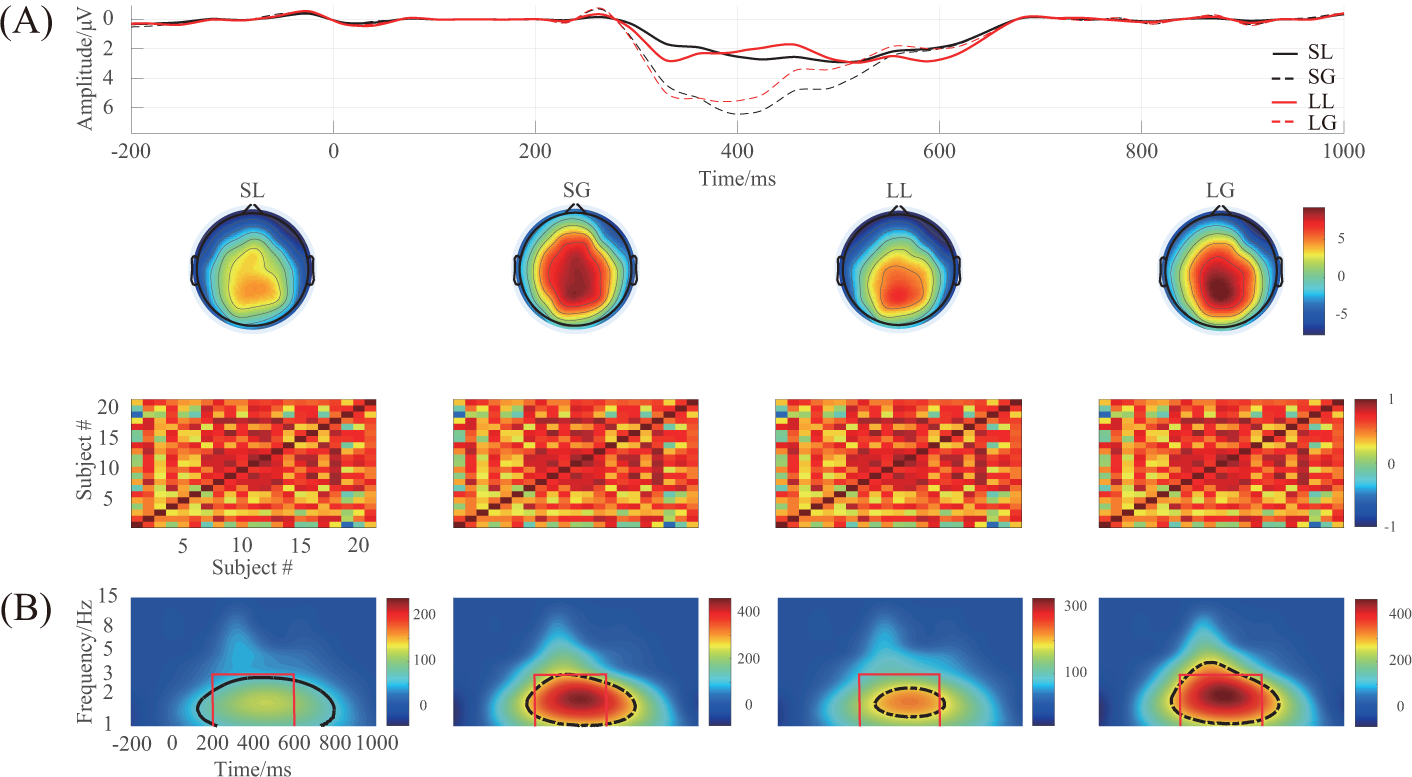
(A) The projected waveform (at Fz, FCz, Cz, CPz, and Pz electrodes), topography (300-600ms), and similarity of topographies across participants of every condition for P3, which were extracted from the real mixed-signal using t-PCA and Promax rotation. (B) The associated grand averaged time-frequency representation (TFR) of every condition for P3-delta oscillation. The red solid rectangle (the time window is 200-600ms and the frequency range is 1-3Hz) for every condition was marked using the conventional rectangle method, the other was gained by the edge detection algorithm.

All displayed topographic maps in the time domain for the simulation and the extracted real datasets were obtained using the peak measurement, and the others were calculated by the mean measurement.

### 2.2. Proposed approach for data processing

In order to overcome the challenges that the EROs cannot be extracted completely via the conventional TFA method because they are affected with each other or artifacts in the time, space, and frequency domains to some degree and the selection of the regions of the associated evoked EROs is subject to the experimenter regarding the time window and the frequency range. The matrix **Z**^*T*^ ∈ ℛ^*M×N*^ (i.e., time samples were variables in columns, and the other factors, such as were integrated into rows which can be labeled as observations) was first formed from the synthetic and real datasets separately to explore the component(s) of interest as the application in the previous investigations (Dien, 2012; Fogarty et al., 2019; Cavanagh et al., 2018; Barry et al., 2019). Next, t-PCA and Promax rotation were fulfilled to decompose this matrix into *R* components, and the components of interest were then selected to project to all of the scalp electrodes to correct their variances and polarities indeterminacy. Subsequently, the calculation of the TFRs of the back-projected components was carried out at some typical electrodes. Finally, the determination of the region of evoked oscillation at the typical electrodes was worked out using the edge detection algorithm for further statistical analysis.

#### 2.2.1. Extracting components of interest and their back-projection

The purpose of the t-PCA and rotation is to use a smaller set of nonredundant descriptive variables (i.e., components) to represent the original ERP signal **Z**^*T*^ and then choose the ones of interest, and the details of the related theories can be seen in Appendix A and Appendix B. Importantly, four steps need to be done during this procedure as below.

The first is about the determination of the number of the remained principal components (PCs). The simple and empirical method, which was to calculate the percentage ratio that the sum of a certain number of lambda values over the sum of all lambda values (each PC has a lambda value), was used to determine the number of the remained PCs (i.e., 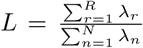, *R* is the number of the retained PCs; *N* is the number of the columns of the matrix **Z**^*T*^, *N ≥ R*; it was named as explained variance of accumulated lambda here) (Bernat et al., 2007; Barry & De Blasio, 2015; Barry et al., 2016; Hu et al., 2015).

The second is about the selection of the rotation method. Dien et al. demonstrated that Promax rotation can generate better results than Varimax rotation (Dien et al., 2005). Additionally, the results of simulation and actual ERP datasets displayed that Promax rotation was more efficient for t-PCA decomposition (Dien et al., 2007). Hence, the Promax rotation was also applied to the study.

The third is about the selection of the temporal and spatial components of interest. If the temporal and spatial properties of the extracted components were consistent with the interested ERP, they will be considered for the next analysis. Overall, in terms of three aspects, the extracted components for the ERP of interest were selected (Barry et al., 2019): (a) The polarity and latency of temporal component; (b) The polarity and location of the excitation region of the spatial component; (c) The similarity of spatial components, herein these are topographies, across all subjects of every condition.

The fourth is about the projection of the selected components to the electrode field. The components, derived from blind source separation algorithms (Comon & Jutten, 2010), herein t-PCA and Promax rotation, had the polarities and the variances indeterminacy, and the back-projection theory can be applied to correct them (Makeig et al., 1997, 1999; Cong et al., 2011a,b). The ERPs were often decomposed into several temporal and spatial components due to the fluctuation of the original waveform of the interested ERP among different subjects. Thus, all of them should be selected to project to the electrode filed to correct their indeterminacy for further analysis.

#### 2.2.2. Transforming the back-projected components into time-frequency representations

For the back-projected data 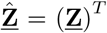 derived from the original signal **Z**^*T*^, we can turn this time domain signal to time-frequency domain signal 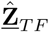 by means of the complex morlet wavelet transformation (Herrmann et al., 2005; Tallon-Baudry et al., 1996; Bertrand & Tallon-Baudry, 2000; Kiebel et al., 2005; Herrmann et al., 2014; Zhang et al., 2020). Specifically, a mother wavelet was first defied using a set of bandwidth and center frequency. Then, the frequency range of interest (e.g., 1-15Hz) and the frequency bins were set for analysis of TFRs.

#### 2.2.3. Objectively determining the region of EROs via edge detection algorithm

The conventional rectangle method was a widely used-way to mark the region of evoked ERO (Ergen et al., 2014; Lally et al., 2014; Jones et al., 2006; Sandre & Weinberg, 2019; Zhang et al., 2019; Her et al., 2019). As the demonstration in Section 3.2.1, different statistical results were displayed because it was a subjective method to determine the region. To address this issue, an edge detection algorithm, Canny detector (Canny, 1986), was used to objectively distinguish the shape of evoked ERO for every condition from TFR distribution. This approach can precisely and objectively mark the position of the oscillatory responses in the TFR distribution based on their shape (time and frequency positions) (Milanović et al., 2019; Saulig et al., 2017; Hory et al., 2002).

This procedure can be divided into the following steps. First, the saved TFR distribution picture of each condition at the specific electrodes was inputted, and the inputted colormap was converted to grayscale. Second, we set the optimal standard deviation, high and low thresholds for the Canny detector. Third, we selected the optimal boundary (as displayed in Figs. 2 - 4, and 6 - 9) for each ERO of interest. Fourth, the value *ψ*_*f,t,c,s*_ of every point (the location of each point is determined by a time point *t* and a frequency bin *f*) of the evoked ERO of interest for *sth* subject under *cth* condition was marked for further statistical analysis, which was performed on the frequency bins, time points, and the pixels of the remained boundary. Last, the mean value 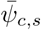of the region of the evoked ERO for every condition of every subject was prepared for investigating the differences among the stimulus types. The details can be seen in Appendix C.

## 3. Results

### 3.1. Synthetic dataset results

During the PCA and Promax rotation procedures, seventeen components were remained, which was explained 99% of variance. According to the temporal and spatial properties of P3 and the similarity of spatial components among all subjects, the 1st, 7th, and 14th components were selected for P3 and they explained 69.85%, 1.60% and 1.20% variance respectively. Likewise, the 2nd and 3rd components were chosen for N2, they explained 6.91% and 2.80% variance respectively.

As shown in Figs. 3 **(B)** and 4 **(B)**, the power of source N2-theta oscillation (about 0.3 *µV* ^2^) was much smaller than that of source P3-theta oscillation (approximately 4 *µV* ^2^) so that the former was easily disappeared in the TFR of the mixed-signal. This was confirmed in the TFR of the mixed-signal as shown in Fig. 3 **(B)** - that is, only one oscillation was observed in the TFR. This was proved by the CC method, specifically, and the CC between the TFRs of the mixed and source/extracted N2-theta was roughly 0.70/0.62 while it was approximately 0.96/0.97 for P3-delta(see Fig. 5, **(G)** and **(I)**). The CCs of the waveforms of the mixed and source/extracted N2/P3 also reflected this from the other side and all of the values were about 0.51/0.96, see Fig. 5 **(A)** and **(C)**. This meant that P3 made the biggest contribution to the mixed-signal which led to the above-mentioned situation and N2 taken account a small part.

Two evoked EROs were obtained corresponding to P3 (see Fig. 3) and N2 (see Fig. 4) respectively using the proposed approach. What is more, the similarity of topographies across all subjects of the extracted signal (especially for N2) was improved using the proposed approach when compared with that of the mixed-signal. Through the comparison of the waveforms/topographies/TFRs of the source, mixed, and extracted signals as shown in Figs. 3 and 4 and the CCs of the these between any two of the three signals (see Fig. 5) for N2-theta/P3-delta, obviously, we found that the CCs of waveforms/topographic distributions/TFRs between the source and the extracted signals for N2/P3 were all roughly equal to 0.99/1.00 (see Fig. 5 **(B), (E)**, and **(H)**), and hence, the conclusion was that the proposed approach can efficiently and objectively extract the ERPs of interest from the mixed-signal which were overlapped in the temporal, spatial, and spectral to some degree which supported the first assumption in the last paragraph of Section 1.

### 3.2. Actual ERP dataset results

#### 3.2.1. Conventional time-frequency analysis method results

Figs. 6 and 7 display the grand averaged waveform, topography, similarity of topographies across all participants, and TFR distribution of each condition for the N2/theta and P3/delta respectively. Tables 1 and 2 were the statistical results of theta and delta oscillations respectively for the three regions determined by the rectangle and the edge detection algorithm.

**Table 1:** The statistical results of N2-theta oscillation for the conventional time-frequency analysis

| Methods | ROI | *WT* |  |  | *FB* |  |  | *WT \* FB* |  |  |
| --- | --- | --- | --- | --- | --- | --- | --- | --- | --- | --- |
| | | *F* | $\eta_p^2$ | *p* | *F* | $\eta_p^2$ | *p* | *F* | $\eta_p^2$ | *p* |
| *CRM* | R1 | 4.198 | 0.173 | 0.054 | 0.293 | 0.014 | 0.594 | 0.032 | 0.002 | 0.860 |
| *CRM* | R2 | 4.278 | 0.176 | 0.052 | 0.692 | 0.033 | 0.415 | 0.003 | <0.001 | 0.959 |
| *EDM* | – | 0.122 | 0.006 | 0.730 | 5.587 | 0.218 | 0.028* | 12.841 | 0.391 | 0.002** |

CRM: the conventional rectangle method; EDM: the edge detection method; \*\* $p < 0.01$ ;
\* $p < 0.05$ ; WT: waiting time; FB: feedback. R1: 100-300ms,3-7Hz; R2: 50-350ms,3-7Hz

**Table 2:** The statistical results of P3-delta oscillation for the conventional time-frequency analysis

| Methods | ROI | *WT* |  |  | *FB* |  |  | *WT \* FB* |  |  |
| --- | --- | --- | --- | --- | --- | --- | --- | --- | --- | --- |
| | | *F* | $\eta_p^2$ | *p* | *F* | $\eta_p^2$ | *p* | *F* | $\eta_p^2$ | *p* |
| *CRM* | R3 | 2.423 | 0.108 | 0.135 | 12.316 | 0.381 | 0.002** | 0.661 | 0.032 | 0.426 |
| *CRM* | R4 | 2.923 | 0.128 | 0.103 | 12.100 | 0.377 | 0.002** | 0.456 | 0.022 | 0.507 |
| *EDM* | – | 0.370 | 0.018 | 0.550 | 7.452 | 0.271 | 0.013* | 1.614 | 0.075 | 0.218 |

CRM: the conventional rectangle method; EDM: the edge detection method; \*\* $p < 0.01$ ;
\* $p < 0.05$ ; WT: waiting time; FB: feedback. R3: 200-600ms,1-3Hz; R4: 300-600ms,1-4Hz

##### N2-theta oscillation

The statistical results of the two regions marked by the conventional rectangle method demonstrated that no significant differences were found for the main and interaction effect respectively as shown in Table 1. However, the interaction effect between the two factors 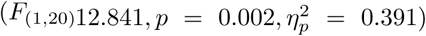 and the main effect of feedback 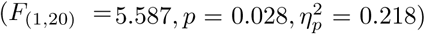 were significant when we used the edge detection algorithm to mark the region of oscillation of interest. The main effect of waiting time did not reach significant level 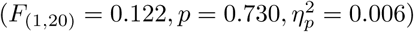.

##### P3-delta oscillation

The statistical results of the edge detection algorithm revealed that the main effect of feedback was significant 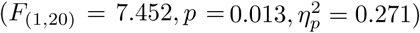 but not for the waiting time 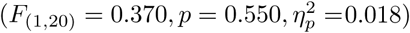. In addition, the interaction effect between waiting time and feedback was also insignificant 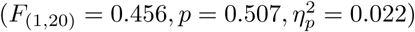. The conventional rectangle results of the two different marked regions also showed the same trends as revealed in Table 2.

#### 3.2.2. Proposed approach results

Figs. 8 and 9 depict the projected waveform, the topographic distribution in the time domain, associated similarity of topographies across all subjects and TFR of every condition for N2-theta and P3-delta respectively. 23 PCs were retained and they accounted for 99% of the variance.

We took the properties of N2 in the temporal and spatial, and the similarity of spatial components across all subjects into account, and the 7th and 9th components were finally selected for next stage and they explained 1.81% and 1.03% of the variance respectively. Note that the polarity of the projected N2 was positive but it can be considered as a negative component, because its original waveform was a negative-going peak (Luck, 2014). For the edge detection algorithm, the statistical results indicated that main effect was insignificant for both waiting time 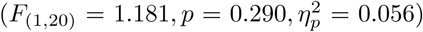 and feedback 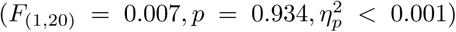. Meanwhile, the interaction effect between waiting time and feedback was also not significant 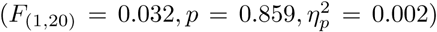. These findings were consistent with our previous study of the results for the time domain analysis (Wang et al., 2014). However, when the conventional rectangle method was performed to determine the region (the time window is 150-300ms; the frequency range is 4-8Hz) for each condition of TFR, we found a significant main effect of waiting time factor 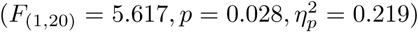 and no other main or interaction effect reached the significant level.

Likewise, for P3, the 1st, 4th, 5th, 11th, 12th, 14th, 15th, and 19th components (explained 56.10%, 4.36%, 3.56%, 0.63%, 0.41%, 0.32%, 0.25%, and 0.17% of the variance respectively) were conducted to project to the electrode field and compute the TFRs via wavelet transform. The statistical results of the edge detection algorithm displayed that there was a significant interaction effect between waiting time and feedback 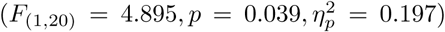.However, no significant interaction effect between them was found for time domain analysis in our previous study (Wang et al., 2014). In addition, the main effects of both waiting time 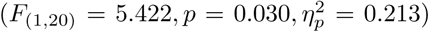 and feedback 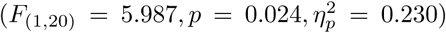 reached a significant level. More concretely, the long waiting time (320.082 ± 74.166 *µV* ^2^) elicited a larger power than that in the short waiting time (256.219 ± 56.869 *µV* ^2^); A large power was observed upon gain condition (346.08 ± 84.335 *µV* ^2^) than lose condition (230.220 ± 48.654 *µV* ^2^), which is similar with the previous investigations (Zhang et al., 2018; Paul et al., 2019; Höltje & Mecklinger, 2020). Then post-hoc analysis was used for further investigation. The results demonstrated that a significant difference was found in the feedback under short waiting time (*p* = 0.011). By contrast, there was an insignificant main effect of feedback under long waiting time condition (*p* = 0.220). When the rectangle method was applied to mark the region (time window is from 200 to 600ms and the frequency range is 1-3Hz), only the significant main effect was obtained for feedback factor 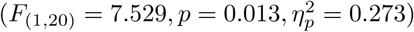.

## 4. Conclusion and discussion

We developed an approach to explore the evoked event-related oscillations (EROs) of interest objectively where (1) Temporal principal component analysis (t-PCA) and Promax rotation were performed to extract the temporal and spatial components of interest simultaneously and project them onto the electrode field to correct their indeterminacies. (2) The time-frequency representations (TFRs) of the projection were computed using the complex morlet continuous wavelet transform. (3) Edge detection algorithm based on the Canny detector was applied on the TFR distributions to recognize the specific time and frequency positions of evoked EROs at some typical electrodes for further statistical analysis.

The yields of the synthetic and actual ERP datasets were governed by the proposed approach better than those of just using the wavelet transform for statistical analysis. Specifically, only one identifiable delta oscillation around 300-600ms can be recognized from the TFR distribution for the mixed synthetic dataset by the conventional time-frequency analysis (TFA) method as shown in the second row in Fig. 3 **(B)**. In contrast, two oscillations were obtained via the proposed approach which corresponded to P3 and N2 respectively as illustrated in Figs. 3 and 4. For the actual dataset P3-delta results, similar to the results of the time domain in the previous reports (Wu & Zhou, 2009; Zhang et al., 2018; Paul et al., 2019; Höltje & Mecklinger, 2020), the loss condition reflected smaller power than by the gain one. Besides, consistent with the findings in the time domain of the paper (Höltje & Mecklinger, 2020)-that is, the interaction effect between the two factors was significant. However, when we applied the conventional TFA method to the signals, only the distinction between the two feedback conditions was found for delta oscillation.

The proposed approach is an efficient and objective method as below. First, the convention TFA method results, which were described in Tables. 1 and 2, revealed that the statistical results of the edge detection algorithm were more stable when compared it with those of the conventional rectangle method which were usually subject to the selected region and were popularly used in previous studies (Ergen et al., 2014; Lally et al., 2014; Jones et al., 2006; Sandre & Weinberg, 2019; Zhang et al., 2019; Her et al., 2019). Second, the statistical results of P3-delta for the conventional TFA method and the rectangle method did not reflect that the feedback was sensitive to the waiting time probably due to the following aspects. (a) The evoked P3-delta was overlapped with other EROs and artifacts to some degree in the temporal, spatial, and spectral, like the TFRs of the conventional TFA method. (b) The information of each ERO of every condition for the projected signal cannot be completely included or some useless information was involved when using the rectangle method to determine the region of P3-delta for each condition. For example, the region was marked as shown in Figs. 8 and 9 **(B)**. However, when we first used the t-PCA and Promax rotation to extract the components of interest, the TFRs of the extracted components was then computed, and the region of ERO of interest was recognized via edge detection algorithm, we found that the power values of feedback for the long and short waiting time were difference which were consistent with the previous study (Höltje & Mecklinger, 2020).

Moreover, the selection of the components in this study also depends on the similarity of the topographies of different subjects and it is expected that different subjects’ topographies are as homogeneous as possible. Regarding one component of the t-PCA, all subjects in one group have the same temporal course, and variant spatial components (i.e., topographies here). This means that, for t-PCA, given an estimated ERP component, the waveform is invariant for all subjects and its topography is variant across all subjects. However, it is strongly expected that the topographies across different subjects for an ERP can be as similar as possible since we expect a homogenous ERP dataset for the repeatable and reliable data analysis. For the results of synthetic and real datasets, the similarities were improved for the projected components to some extent when these were compared with those of the mixed-signal (especially for N2 of the extracted signal as illustrated in the last column the Fig. 4). This demonstrated that the homogeneity of the topographies of different subjects was better than before with the proposed approach.

The future investigation can be carried out in the following aspects. First, the evoked oscillations extracted from the ERP waveforms using the proposed approach are time-locked and phase-locked but some important information like induced oscillation is neglected. The induced oscillation (Tallon-Baudry et al., 1996) is canceled out by the averaging procedure over trials in the time domain (Tallon-Baudry & Bertrand, 1999; Pfurtscheller & Da Silva, 1999). Additionally, induced oscillation is generated by nonlinear and possibly autonomous mechanisms, and high-order processes whereas evoked oscillation is related to stimulus-locked (David et al., 2006). Then, it is extremely used to investigate the neural mechanisms, such as attention modulation (Herrmann & Knight, 2001), the gamma brain oscillatory in healthy participates, and cognitive impairment patients (Başar, 2013) and so on. Therefore, it is interesting to use our method to extract induced oscillatory responses in the future. Second, some TFA techniques, like the combination of Wigner-Ville distribution and Gabor transform with the matching pursuit decomposition which can provide an appropriate time-frequency resolution in all frequencies (Wacker & Witte, 2013), can be also attempted to extract evoked and induced oscillations.

## Acknowledgments

This work was supported by the National Natural Science Foundation of China (Grant No.91748105) and the Fundamental Research Funds for the Central Universities [DUT2019] in Dalian University of Technology in China, and the scholarships from China Scholarship Council (No.201806060165). The authors also would like to thank Professor Peng Li who works at Shenzhen University for sharing their ERP datasets with us in this study. Thanks for Guoqiang Hu, Xichu Zhu, and Xueyan Li helping us to improve the quality of the draft.

## Appendix A. The explanation of temporal-PCA and Promax rotation from the view of blind source separation

When the temporal-PCA and Promax rotation (Hendrickson & White, 1964) are applied to decompose an ERP dataset **Z** ∈ ℛ^*N×M*^ (N and M respectively represent the number of time points within one epoch and the number of sensors of all subjects under all conditions) which is collected from brain scalp of multi-subject and multi-condition and these can be interpreted via the linear model (Cong et al., 2011a,b).

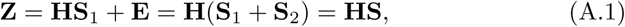

where, **H** is the mixing matrix with full rank; **S** = **S**_1_ + **S**_2_ (**S** ∈ ℛ^*R×M*^), **E** = **HS**_2_, they are the unknown correlated source signals and the sensor noise respectively.

As described in (Cong et al., 2011a), the assumption of the model in Eq. A.1 is over-determined and it means that the number of the observed signals *N* is larger than that of the source signals *R*. Once the estimation of the number of the sources is done (here, the determination is based on the explained variance of accumulated lambda as mentioned in Section 2.2.1 in this study), the overdetermined model can be changed to the determined one as follows

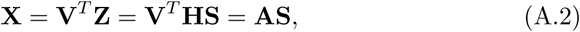

where **X** ∈ ℛ^*R×M*^; **V**^*T*^ ∈ ℛ^*R×N*^ represents the dimension reduction matrix generated from performing PCA on **Z**^*T*^ ; **A** = **V**^*T*^ **H, A** ∈ ℛ^*R×R*^ is also named the mixing matrix.

Aiming to separate the mixture in Eq. A.2, the blind source separation algorithm can be applied (Comon & Jutten, 2010). Here, Promax rotation (Hendrickson & White, 1964) is used to obtain an unmixing matrix **W** in this study. The algorithm of Promax rotation is described in Appendix B and the inverse matrix **B** = **W**^−1^ represents the estimation of **A** (Cong et al., 2011a).

And then we use this unmixing matrix **W** to turn the mixture in Eq. A.2 into a matrix of estimated components as below (Cong et al., 2011a).

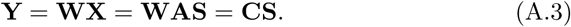

In the above formulation, **Y** is the estimation of **S** and its each row can be assembled to the scalp-map/topography of multi-subject of multi-condition; **C** = **WA** is the global matrix.

Under this determined model condition in Eq. A.2, aiming to solve the issue that different components, which is derived from the matrix **X**, cannot be statistically further analyzed together in terms of peak amplitudes because their polarities and amplitudes are indeterminate, we project the component(s) of interest back to the electrode field in this study which is often used in many other blind source separation for EEG, for example, independent component analysis procedures (Makeig et al., 1997, 1999; Cong et al., 2011a,b). The back-projection can be illustrated as (Cong et al., 2011a,b).

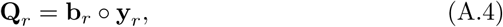

here, **Q**_*r*_ ∈ ℛ^*R×M*^ is the projection of *rth* component at all electrodes, **b**_*r*_ is the *rth* column of the inverse matrix **B**, and **y**_*r*_ is the *rth* row of the estimated matrix **Y**.

Under the global optimization, there exists only one nonzero element in each row and each column of **C**. In other words, the *rth* extracted component is unknown scaled version of the *mth* source signal. Hence, the projection in Eq. A.4 can be described as (Cong et al., 2011a,b)

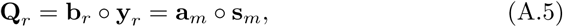

where, **a**_*m*_ is the *mth* column of the mixing matrix **A**, and **s**_*m*_ is the *mth* row of the source matrix **S** in Eq. A.2.

To the original overdetermined model in Eq. A.1, the *rth* component derived from the matrix **Z** can be projected to the all electrodes as below

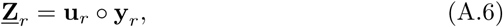

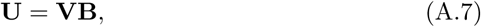

where **u**_*r*_ is the *rth* column of **U** and denotes the time-course/waveform. We use combination of the inverse matrix **B** and the dimension reduction matrix **V** to represent **U**, which is achieved to estimate the mixing matrix **H** in Eq. A.1 (Cong et al., 2011a).

In most cases, several components need to be projected back to all electrodes simultaneously (Cong et al., 2011a,b; Delorme & Makeig, 2004), hence, the related projection of several components can be implemented as follows

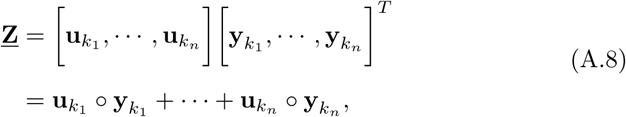

where *k*_1_, *…, k*_*n*_ (1 *≤ k*_*n*_ < *R*) are the number of selected components; ‘*°*’ denotes the outer product of vectors. The size of each dimension of the matrix **<u>Z</u>** is the same with that of **Z**.

## Appendix B. Oblique Procrustes transformation

Mathematically, Procrustes equation can be defied as (Hendrickson & White, 1964; Richman, 1986):

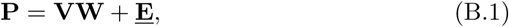

where, **P** is the pattern matrix; **W** is the transformation matrix; **<u>E</u>** is the residuals matrix. The satisfactory result is that we can find a transformation matrix to make the value of **<u>E</u>**^*T*^ **<u>E</u>** as close zero as possible.

Specifically, the pattern matrix **P** is first generated from the matrix of un-rotated factor loadings **V** by the target Procrustes transformation, and the determination of **V** is based on PCA and ‘factor’ is also called as principal component here.

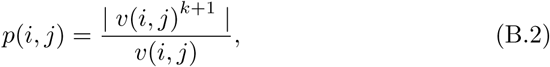

where, *k >* 1. The matrix **P** stands for the matrix **V** raised to the *kth* power and its original sign is unchanged.

Next, the least squares method is performed to calculate the fit of the orthogonal matrix **V** of the factor loadings to the pattern matrix so that **<u>E</u>**^*T*^ **<u>E</u>** is a minimum.

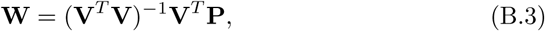

where **W** is the transformation matrix in Promax rotation (Richman, 1986); **V**^*T*^ is the transpose matrix of the orthogonal rotated matrix **V**; (•)^−1^ is the inverse of a matrix.

## Appendix C. Determination of the region of Evoked ERO via Canny detection algorithm

As shown in Fig.5, the interested ERO of each condition has a boundary that clearly distinguishes it from others in the grand averaged TFR distribution (this is usually generated from 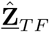 by calculating the mean values of the specific electrodes, we use the symbol *φ*_*f,t,c,s*_ to represent the value of any point in the TFR distribution for *s*^*th*^ subject under*c*^*th*^ condition) for EEG signal. Following this context, we can use a typical approach, canny detection algorithm, to determine the optimal boundary and then gain the associated region of ERO.

The procedure of the original Canny algorithm for the determination of the boundary of a target can be approximately divided into the following steps (Canny, 1986; Deng et al., 2013; Xu et al., 2014).

First, any noise is filtered out from the original image using a Gaussian filter (it is computed by a simple mask so that it is widely used in the Canny algorithm) before trying to use this detector to detect any edges. Actually, this step is to calculate the convolution between the raw image and the mask.

Second, calculation of the gradient amplitude and direction at any pixel location, which is aim to find the edge strength. The gradient amplitude is determined as the square root of the sum of the square of the horizontal *G*_*x*_(*i, j*) and vertical gradient *G*_*y*_(*i, j*) amplitudes.

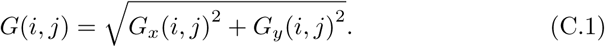

Then,the gradient direction at every pixel can be defined as:

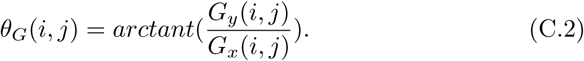

Third, the non-maxima suppression is applied to the gradient amplitude to make the blurred edges become sharper. In other words, the gradient direction at every pixel is computed to find the maximum magnitude. For one thing, when the gradient direction of this pixel is considered as one of 8 possible primary directions (i.e., 0*°*, 45*°*, 90*°*, 135*°*, 180*°*, 225*°*, 270*°*, and 315*°*), the comparisons are made between the gradient magnitude of this pixel and its two neighbors along the gradient direction. If this value is the greatest one, then it is remained, and otherwise it will be set to zero. For another thing, if the gradient direction is not belonging to any of these possible directions, it can be finished to calculate the neighboring gradients based on interpolation theory (Xu et al., 2014).

Fourth, the determination of edge map via hysteresis thresholding. It needs two thresholds to better recognize the edges: a high threshold *T*_1_ and a low one *T*_2_. If any pixels have the values (i.e., the gradient magnitude G(i,j)) greater than *T*_1_ are looked as strong edges, and then it will be recorded. Meanwhile, the gradient amplitudes of the pixels are greater than *T*_2_ and are connected to the strong edges, they will be selected as a strong one. Otherwise, they are not included in the final edge image.

Practically, the region of interest needs to be determined based on the recognized boundary for further statistical analysis. Any position (it is determined by a frequency bin-*f*_1_ and a time point-*t*_1_) within the marked boundary is first calculated by performing on the frequency bins, time points, and the pixels of the boundary. Each value *ψ*_*f,t,c,s*_ of the point of the related ERO is following remained for every subject *s* under each condition *c* at the channels of interest.

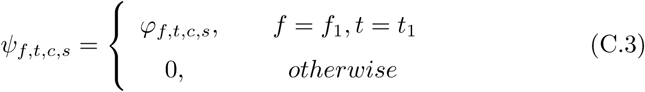

Last, the demanded value 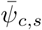 for each subject of every condition is gained by computing the mean value of the marked evoked ERO.

